# Frequency-specific maturation of cortical organization and social-cognitive links from childhood to adolescence

**DOI:** 10.1101/2025.09.01.673446

**Authors:** Zhu-Qing Gong, Elizabeth Jefferies, Xi-Nian Zuo

## Abstract

The human brain unfolds its functional architecture over multiple timescales during childhood and adolescence, yet we know little about how these developmental trajectories vary between individuals. Slow brain signals at different frequencies are thought to support distinct functional processes, but their contribution to shaping large-scale cortical organization across development remains unclear. Here, we map the maturation of cortical organization by decomposing spontaneous slow brain activity into multiple frequency bands in participants aged 6-19 years. We identify a reproducible three-stage progression – child-like, transitional, and adult-like configurations – whose timing depends on frequency: faster dynamics mature earliest, intermediate frequencies develop gradually, and lower-frequency dynamics reorganize abruptly around puberty. Individual cognitive and social traits modulate these trajectories: children with higher social anxiety or higher IQ reach adult-like configurations earlier in specific frequency bands, whereas lower-IQ children show generalized delays. These results reveal that cortical organization matures along multiple, frequency-specific timescales and that intellectual and socio-emotional factors shape its pace. Our multi-frequency framework provides a new perspective on hierarchical brain development and may inform biomarkers for atypical neurodevelopment.

## Introduction

Human cortical organization is captured by connectivity gradients running from sensorimotor to association cortex (Margulies et al., 2016). This cortical hierarchy emerges early in development: by 25 weeks of gestation, the fetal brain already exhibits gradient-like patterns within resting-state networks, with primary sensory and motor systems showing consistent organization while higher-order networks are more variable (Moore et al., 2023). During childhood and adolescence, these large-scale gradients become increasingly refined, paralleling behavioral maturation. The principal gradient, initially anchored between sensorimotor and visual regions, rotates into an adult-like configuration by mid-adolescence, separating transmodal default-mode and control networks from unimodal areas, with rapid change around 12 years (Dong et al., 2021; Xia et al., 2022). The ventral attention network appears to modulate this process, with lower connectivity in this network linked to accelerated maturation and improved cognitive performance (Dong et al., 2024). Whole-brain analyses further show that this sensorimotor-to-association axis persists across the lifespan, with early childhood changes reinforcing hierarchical structure and later trajectories reflecting system-specific maturation and decline (A. C. Luo et al., 2024; Sun et al., 2025). Collectively, these findings highlight adolescence as a pivotal period for cortical reorganization (Dahl, 2004; Steinberg, 2005), yet the temporal dynamics and individual variability of these changes remain poorly understood.

Elucidating how this cortical reconfiguration tracks cognitive and social growth is essential for understanding the neural basis of mental development during this formative period. The present study addresses this goal by examining these developmental processes across distinct frequency bands. Crucially, brain functional activity is intrinsically dynamic across multiple temporal scales (Buzsaki & Draguhn, 2004; Buzsaki & Voroslakos, 2023; Klimesch, 2018). This frequency-dependent property of neural dynamics holds across measurement modalities, including blood-oxygen-level-dependent (BOLD) signals acquired with functional MRI (fMRI). However, early fMRI studies, constrained by hardware and sequence limitations, focused on a narrow low-frequency range (∼0.01-0.1 Hz), a convention that has largely persisted. Recent advances in scanner performance, fast multiband acquisition, and improved denoising strategies (e.g., ICA-based approaches rather than strict bandpass filtering) have expanded the usable frequency range of BOLD signals, enabling multi-band analyses that more fully probe frequency-specific neural properties. High-temporal-resolution multiband acquisitions can capture BOLD oscillations spanning six slow-frequency bands (slow-1 through slow-6) (Buzsaki & Draguhn, 2004; Penttonen & Buzsáki, 2003). Ongoing BOLD signals indeed fluctuate across these bands, each potentially indexing distinct neurophysiological processes and functional properties (Gong & Zuo, 2024; Penttonen & Buzsáki, 2003; Ries et al., 2018; Zuo et al., 2010). A growing multi-band fMRI literature has begun to reveal such frequency-specific signatures, yet the developmental trajectories of these frequency-resolved signals remain largely unexplored.

Intrinsic fMRI activity reflects a multilayered system of neural oscillations that support brain function even without tasks (Raichle, 2006; Fox & Raichle, 2007; Tavor et al., 2016). Building on 15 years of multiband fMRI, we proposed a hierarchical framework describing how distinct frequency bands jointly contribute to this system (Gong & Zuo, 2025). Higher frequencies (slow-1 to slow-3, roughly 0.08–1.64 Hz) have been linked to detection and rapid perceptual processing, intermediate bands (slow-4, ∼0.03–0.08 Hz) to relatively segregated cognitive operations, and lower bands (slow-5 and slow-6, ∼0.004–0.03 Hz) to long-range integrative functions. This organization mirrors evidence from electrophysiology, where fast oscillations support local processing and slower rhythms support distributed associative functions (Buzsaki & Draguhn, 2004; Engel et al., 2001; Kopell et al., 2000). Anatomically, this frequency differentiation aligns with axonal properties—local sensory and motor areas are connected by thick, heavily myelinated fibers conducting fast signals, whereas distant association cortices are linked by thinner, slower-conducting fibers (Aboitiz, 1992). These shared structural constraints help explain why neural activity recorded with different modalities (e.g., EEG, MEG, fMRI) exhibits broadly comparable frequency architecture, consistent with the fractal nature of neural dynamics: across measurement scales, from single-unit recordings to macroscopic EEG and fMRI, frequency structure shows principled similarity.

Empirical studies increasingly suggest that distinct BOLD frequency bands support different functional domains and cognitive states. For example, specific bands have been linked to particular cognitive processes (Gong & Zuo, 2024) and to varying functional segregation and integration patterns (Ma et al., 2021; Park et al., 2019; Thompson & Fransson, 2015). Although the overall spatial architecture of cortical gradients is preserved across bands, individual networks reach peak integration at distinct frequency ranges (Gong & Zuo, 2023), showing that resting-state networks operate across multiple intrinsic tempos (Gohel & Biswal, 2015; Z. Luo et al., 2024; Nese et al., 2024; Zhang et al., 2015). Yet little is known about how the developmental refinement of cortical gradients depends on the timescales of neural oscillations. Most developmental studies have focused only on conventional low frequencies (∼0.01–0.1 Hz, slow-4 and slow-5), potentially missing band-specific maturation. Disentangling BOLD signals across bands allows us to test whether hierarchical reorganization follows distinct trajectories at different tempos of brain activity.

Another open question is how individual differences in cognition and social experience influence the maturation of functional connectivity gradients. Childhood and adolescence show wide variability in intelligence and sensitivity to the social environment (Haworth et al., 2010; Venticinque et al., 2024), and developmental changes in neural oscillations may contribute to these differences (Cellier et al., 2021; Easson & McIntosh, 2019; Rier et al., 2024; Tan et al., 2024; Turri et al., 2023). For example, steeper age-related increases in task-evoked frontoparietal theta power predict higher working memory and fluid intelligence scores (Tan et al., 2024), and reading skill relates to connectivity between networks (Lex et al., 2025). High-IQ children also show greater reorganization of network topology, with stronger integration, segregation, and homotopic symmetry (Suprano et al., 2019), suggesting that advanced cognition depends on efficient large-scale network coordination. Social development shows a similar pattern: adolescence, a period of heightened peer sensitivity, coincides with major network reorganization (Crone & Dahl, 2012; Somerville, 2013). Children especially attuned or reactive to social contexts might exhibit precocious functional maturation of “social brain” networks (Kirby et al., 2018; Skyberg et al., 2023), while chronic social stress is linked to atypical or delayed trajectories (Bick & Nelson, 2016; Callaghan & Tottenham, 2016; Tooley et al., 2021). Despite these insights, it remains unknown whether multi-band functional gradients follow distinct developmental trajectories in children with different intellectual capacities or social sensitivities, even though different BOLD frequency bands have distinct functional profiles. Conceptually, multi-band analysis simply partitions the same BOLD signal, with single-band approaches as a special case; and in practice, harmonized preprocessing with ICA-based denoising (e.g., ICA-AROMA) further limits vascular or developmental confounds. Addressing this gap may reveal how cognitive and social factors shape the timing and organization of large-scale functional architecture in the developing brain.

In this study, we systematically examined the development of resting-state functional connectivity gradients in children aged 6–19 across multiple BOLD frequency bands. We tested whether large-scale network integration emerges at different ages depending on frequency, by decomposing each child’s BOLD time series into distinct bands and computing cortical connectivity gradients for each age group. We also asked whether developmental trajectories vary with intellectual ability or social anxiety, probing whether advanced cognition or heightened social-affective engagement is linked to accelerated maturation of functional gradients. This multi-band, multi-dimensional approach builds on recent advances in connectome gradient mapping and highlights the value of considering neural oscillation spectra in neurodevelopment. Together, our findings show how frequency-specific dynamics shape large-scale brain organization, and how individual differences in cognition and social experience modulate the timeline of network integration.

## Results

### Developmental trajectories of the first two functional connectivity gradients across frequency bands

The frequency range and the number of decomposable BOLD bands are determined by the fMRI sampling rate and scan duration. We decomposed the preprocessed BOLD time series using the DREAM toolkit, which partitions the signal into slow-frequency bands on the basis of the N3L framework given the acquisition parameters (Gong et al., 2021). Restricted by our fMRI sampling parameters, the spectrum supported three low-frequency bands: slow-3 (0.0822–0.2000 Hz), slow-4 (0.0311–0.0822 Hz), and slow-5 (0.0133–0.0311 Hz). We divided the full cohort (ages 6–19 years) into twelve age groups (combining ages 6–7 and 18–19 into single groups due to low sample sizes at these extreme ages, Fig. 1A). Because the first and second gradients capture a substantial proportion of the variance in functional connectivity and can be assigned relatively clear physiological interpretations, our analyses concentrated on characterizing their developmental trajectories across multiple frequency bands. Consistent with prior single-band findings, the first two gradients exhibited a developmental “flip” or reversal during adolescence (Dong et al., 2021; Xia et al., 2022). However, our frequency-resolved approach revealed a more refined picture: the maturation of these gradients unfolded in a reproducible three-stage trajectory, rather than two, across all bands (Fig. 1B). Although the age spans of these stages differed by band, their cortical topographies were similar across bands. In Stage I, the first gradient resembled the adult second gradient, characterized by a sensory-motor axis extending from visual cortex through auditory areas to somatomotor regions. Conversely, the second gradient at this stage mirrored the adult first gradient, capturing the primary-to-association cortical hierarchy. Stage II was marked by a transitional pattern in which sensorimotor and visual regions became increasingly separated, while the default mode network (DMN) emerged at one extreme of the gradient axis. By Stage III, both gradients converged onto adult-like configurations: Gradient 1 was dominated by the unimodal-to-transmodal hierarchical axis, and Gradient 2 by the differentiation among sensory and motor systems. This three-stage progression suggests a temporally phased refinement of cortical organization, with each stage defined by distinct, though gradually evolving, gradient topographies.

**Fig. 1.**
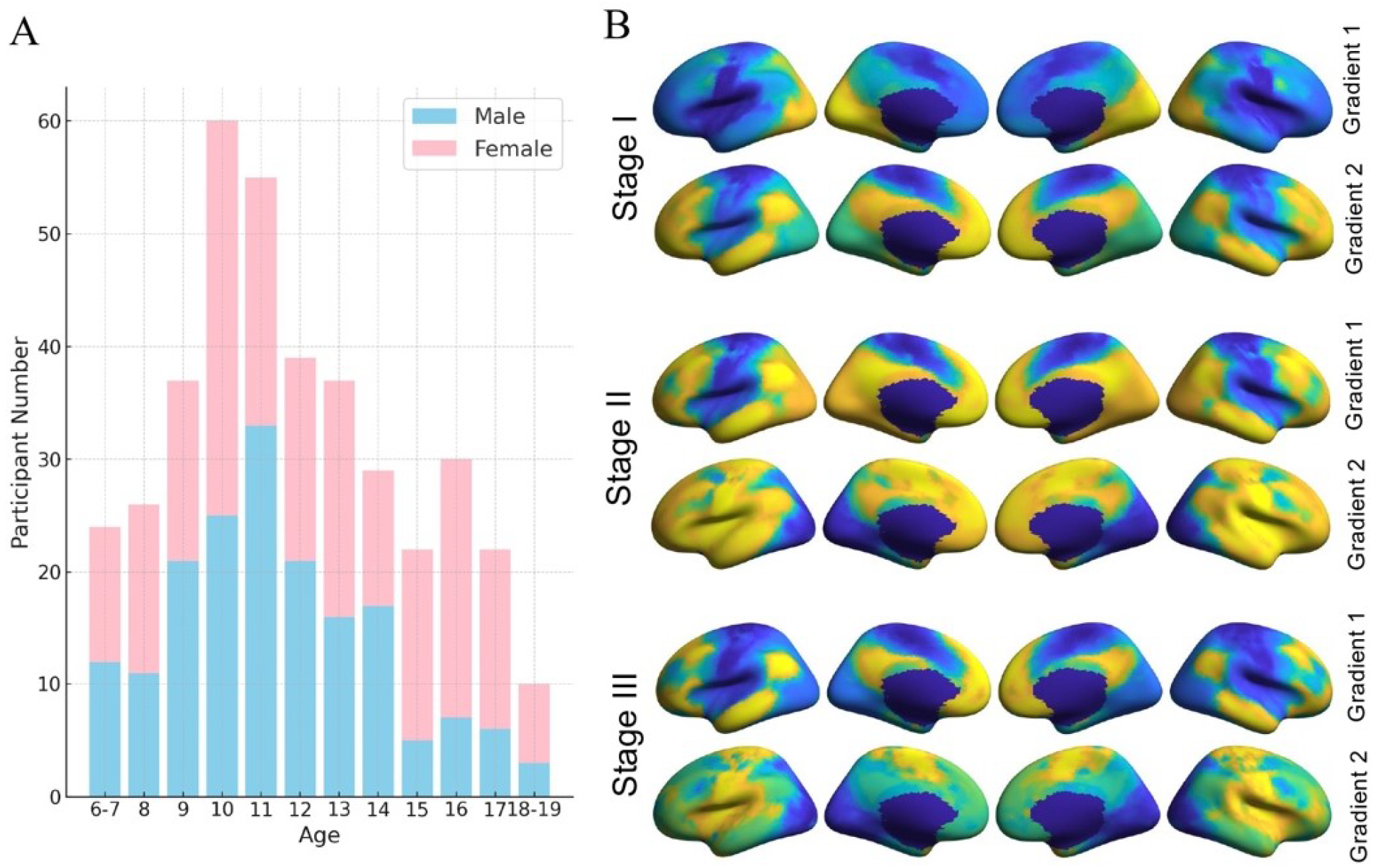
(A) Participants were divided into 12 age groups. The x-axis denotes age groups; the y-axis indicates the number of participants per group. Blue bars represent boys, and pink bars represent girls. (B) Three-stage developmental trajectories of multi-band functional connectivity gradients during the school-age period. In Stage I, Gradient 1 reflects the segregation of primary sensorimotor regions, while Gradient 2 captures a hierarchical axis from primary to association areas. In Stage II, both gradients exhibit a mixed configuration, with visual and somatosensory regions occupying opposite ends of the gradient axis and the default mode network positioned variably at either end. In Stage III, the gradients reverse relative to Stage I, forming an adult-like configuration.

We then assigned, for each band and age group, the group-level gradient maps to one of these stages. To preserve developmental continuity and ordering, occasional single-age “back-and-forth” assignments were aligned with adjacent age groups, yielding a single monotonic pass through Stages I–III with increasing age in each band. The timing of these gradient transitions differed markedly across frequency bands (Fig. 2, Fig. S1–S3). For the first gradient (Fig. 2A), the onset of Stage II (the transition from child-like to intermediate pattern) and Stage III (arrival at adult-like pattern) occurred at younger ages in the higher-frequency band and at later ages in the lower-frequency bands. In slow-3, Stage II began at ∼9 years and Stage III at ∼12 years. In slow-4, Stage II emerged by ∼6–7 years, but Stage III did not appear until ∼14 years. In slow-5, Stage II began around ∼12 years, with Stage III following by ∼13 years. For the second gradient (Fig. 2B), maturation lagged behind the first gradient in all bands. Slow-3 showed Stage II and III transitions at ∼9 and ∼16 years, respectively. In slow-4, Gradient 2 entered Stage II by ∼6–7 years but remained in that transitional state through late adolescence, achieving Stage III only at ∼18–19 years. In slow-5, Gradient 2 underwent its Stage II and III shifts around ∼12 and ∼13 years, paralleling the rapid timeline of Gradient 1 in that band. These differential maturation rates suggest a sequential refinement process in brain development: large-scale functional integration (captured by Gradient 1’s hierarchy) is achieved earlier, followed by a more protracted segregation or differentiation of functional domains (captured by Gradient 2). This pattern was most pronounced in the slow-4 band, which in our framework corresponds to the intermediate “computation” level supporting foundational cognitive and motor functions, indicating that although these networks begin maturing in early childhood, their refinement extends over a longer developmental window. Summarizing across bands, slow-3 reached adult-like organization earliest, slow-5 showed a compressed peri-pubertal shift, and slow-4 exhibited the earliest onset but longest duration. Notably, the peri-pubertal flip in slow-5 aligns with timings reported in single-band (0.01–0.1 Hz) analyses, indicating that single-band transitions are likely driven primarily by slow-5 dynamics.

**Fig. 2.**
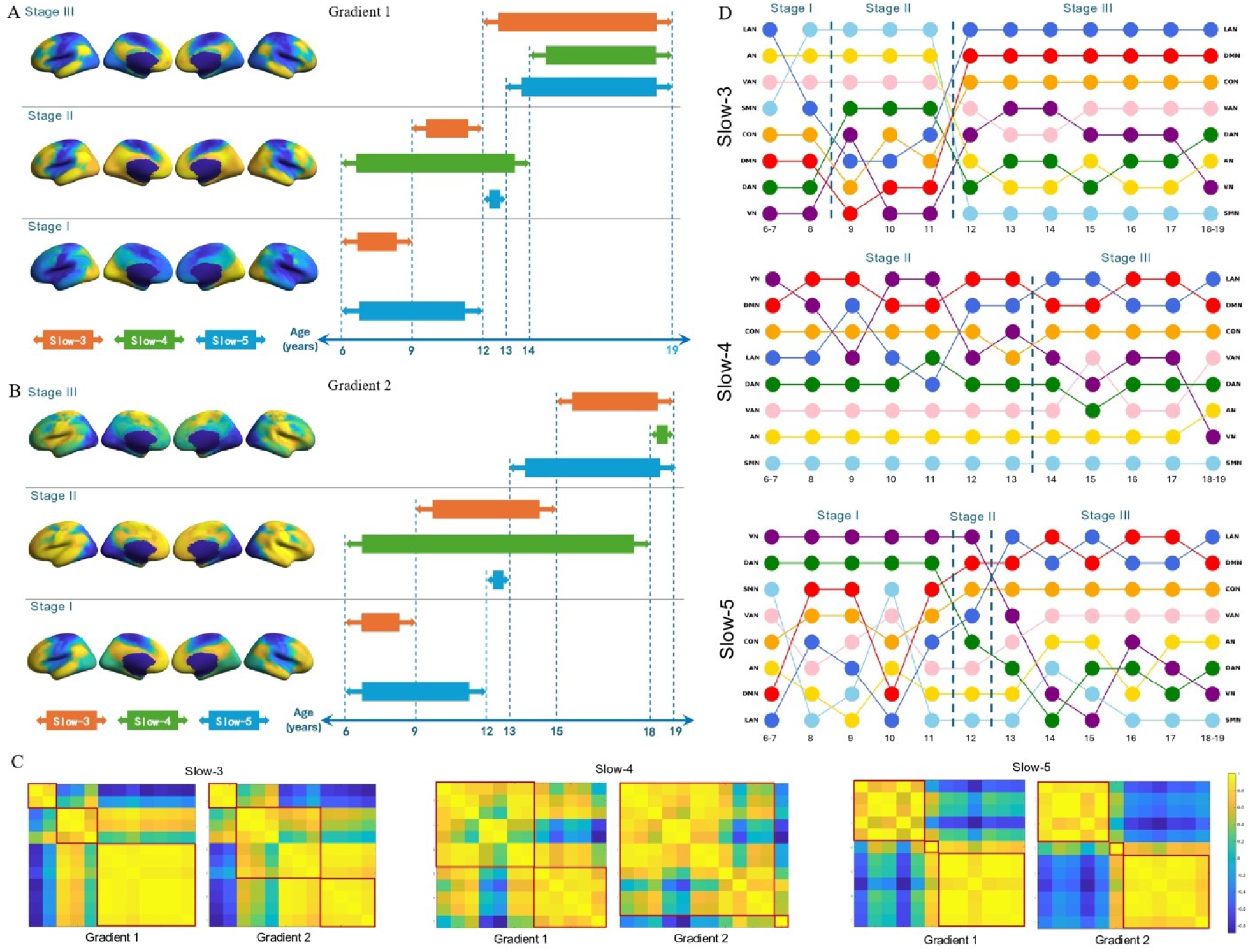
(A, B) Developmental timelines of the first and second functional connectivity (FC) gradients across three frequency bands. The left panels indicate the three developmental stages (Stages I–III) for Gradient 1 (A) and Gradient 2 (B), respectively. The right panels show the age range during which each frequency band remains in a given stage. (C) Pairwise spatial correlations of gradient maps across all age groups within each frequency band. Significance was assessed with a 1,000-iteration spin permutation test. Both axes represent the 12 age groups, ordered from youngest (bottom/left) to oldest (top/right). Red boxes highlight within-stage correlations. Overall, within-stage correlations were robustly positive across bands. Notably, Stage II exhibits greater within-stage heterogeneity compared to Stages I and III. (D) Age-related shifts in the positions of eight canonical large-scale brain networks along the first FC gradient, shown separately for each BOLD frequency band. Each column represents an age group, and each color corresponds to a specific functional network.

Because our staging was derived from observed maturational patterns in the gradient maps, we next quantified similarity within and across stages by computing pairwise spatial correlations between all age groups for each frequency band and gradient (Fig. 2C). Significance was assessed with a 1,000-iteration spin permutation test. Overall, within-stage correlations were robustly positive across bands: Stages I and III showed consistently high similarity (median r ≳0.90 for both gradients), whereas Stage II was lower and more variable (typically r ≈0.5–0.9) with a small number of age-pairs not reaching significance. Specifically, for the slow-3 band, both gradients exhibited strong within-stage similarity in Stages I and III, with greater dispersion in Stage II; the slow-4 band showed a prolonged Stage II (especially for Gradient 2) with broadly significant but more heterogeneous correlations; and in slow-5, the brief Stage II and rapid consolidation into Stage III were reflected in uniformly high within-stage similarity thereafter. Together, these patterns validate the three-stage segmentation and indicate that Stage II captures a transitional window marked by heightened heterogeneity, consistent with individual differences in developmental pacing and potentially elevated neuroplasticity in early adolescence.

Fig. 2D illustrates how the eight canonical large-scale networks shift along the principal connectivity gradient as a function of age in each BOLD frequency band. Although the broad three-stage topographies are conserved across bands, the developmental choreography of individual networks is markedly frequency specific. In the slow-3 band, the language network undergoes the most pronounced ascent toward the transmodal apex during childhood, after which it stabilizes—together with the default-mode, control, and somatomotor networks—once Stage III is reached at approximately age 12. By contrast, the two attention networks and the unimodal visual and auditory networks continue to exhibit modest fine-tuning, indicating ongoing optimization of sensory-attention coupling even after the major reorganization is complete. The slow-4 band reveals a different pattern. Here, somatomotor and auditory networks achieve near-adult positions as early as 6–7 years and remain stable throughout the school-age interval, whereas the visual network shows the most protracted trajectory, descending steadily toward the unimodal pole. After entry into Stage III at about 14 years, residual changes are largely confined to continuous changes of visual network, subtle reciprocal refinements between language and default-mode networks, and between ventral and dorsal attention networks. In the slow-5 band, the two attention networks follow opposite courses: the ventral network continues to shift until mid-adolescence before plateauing, whereas the dorsal network remains static in childhood but begins a sustained downward movement around puberty. Most large-scale realignments in this band are compressed into the peri-pubertal window; after the brief transitional Stage II centred on age 12, later adolescence is dominated by local adjustments between language and default-mode networks and between dorsal attention and sensory and motor networks. Taken together, despite sharing a common three-stage framework, each frequency band shows a distinct timetable and magnitude of network reordering: slow-3 achieves early stabilization of high-level networks, slow-4 extends the visual-driven refinement window, and slow-5 concentrates its major shifts around puberty. The results reinforce the notion that cortical maturation unfolds on multiple temporal scales, each preferentially expressed in a distinct oscillatory band and reflected in the frequency-specific reordering of large-scale networks along the cortical hierarchy.

### Quantifying rates and modes of first-gradient development across frequency bands

To further quantify between-band differences in developmental rate and mode, we derived three subject-level metrics from the spatial distribution of the first gradient: (i) the first-gradient integration score (FGIS), indexing the degree to which the principal gradient expresses a unimodal-to-transmodal integration axis; (ii) the first-gradient segregation score (FGSS), indexing segregation of sensory-motor systems; and (iii) the first-gradient integration-segregation score (FG-ISS), a signed composite where higher (positive) values indicate an integration-dominant configuration (Stage III), negative values indicate a segregation-dominant configuration (Stage I), and values near zero reflect a mixed, transitional configuration (Stage II). Using 1–3 waves of longitudinal data per child, we modeled age-related trajectories of FG-ISS, FGIS, and FGSS in each band with generalized additive mixed models (GAMMs).

FG-ISS trajectories (Fig. 3A) showed robust age effects in all bands (P<0.001), with explained variance R^2^: slow-5 = 0.454 > slow-3 = 0.356 > slow-4 = 0.336. All three bands displayed nonlinear increases with age (faster gains in childhood, tapering in adolescence). Notably, FG-ISS values of slow-5 were below zero before ∼13 years and above zero thereafter, indicating a shift from segregation-dominant (Stage I) to integration-dominant (Stage III) organization around early adolescence, consistent with the three-stage model. The slow-4 curve hovered near zero until ∼14 years, mirroring the prolonged Stage II in this band. Slow-3 started with a lower intercept but a steeper childhood slope, suggesting greater early segregation yet the fastest developmental rate among bands. Integration-specific indices (Fig. 3B) reinforced this pattern: age effects were significant across bands (P<0.001; R^2^: slow-5 = 0.373 > slow-3 = 0.309 > slow-4 = 0.055). FGIS increased with age in slow-5 and slow-3, whereas slow-4 began at a comparatively high integration level and remained nearly flat across the school-age years, consistent with integration in this band having already advanced by pre-school age. Segregation-specific indices (Fig. 3C) also showed significant age effects in all bands (P<0.001; R^2^: slow-5 = 0.379 ≈ slow-4 = 0.373 ≈ slow-3 = 0.308), with FGSS declining with age. Taken together, the combination of rising FG-ISS and falling FGSS indicates that slow-4 reorganization is driven primarily by decreasing segregation, whereas slow-3/slow-5 show both increasing integration and declining segregation across development.

**Fig. 3.**
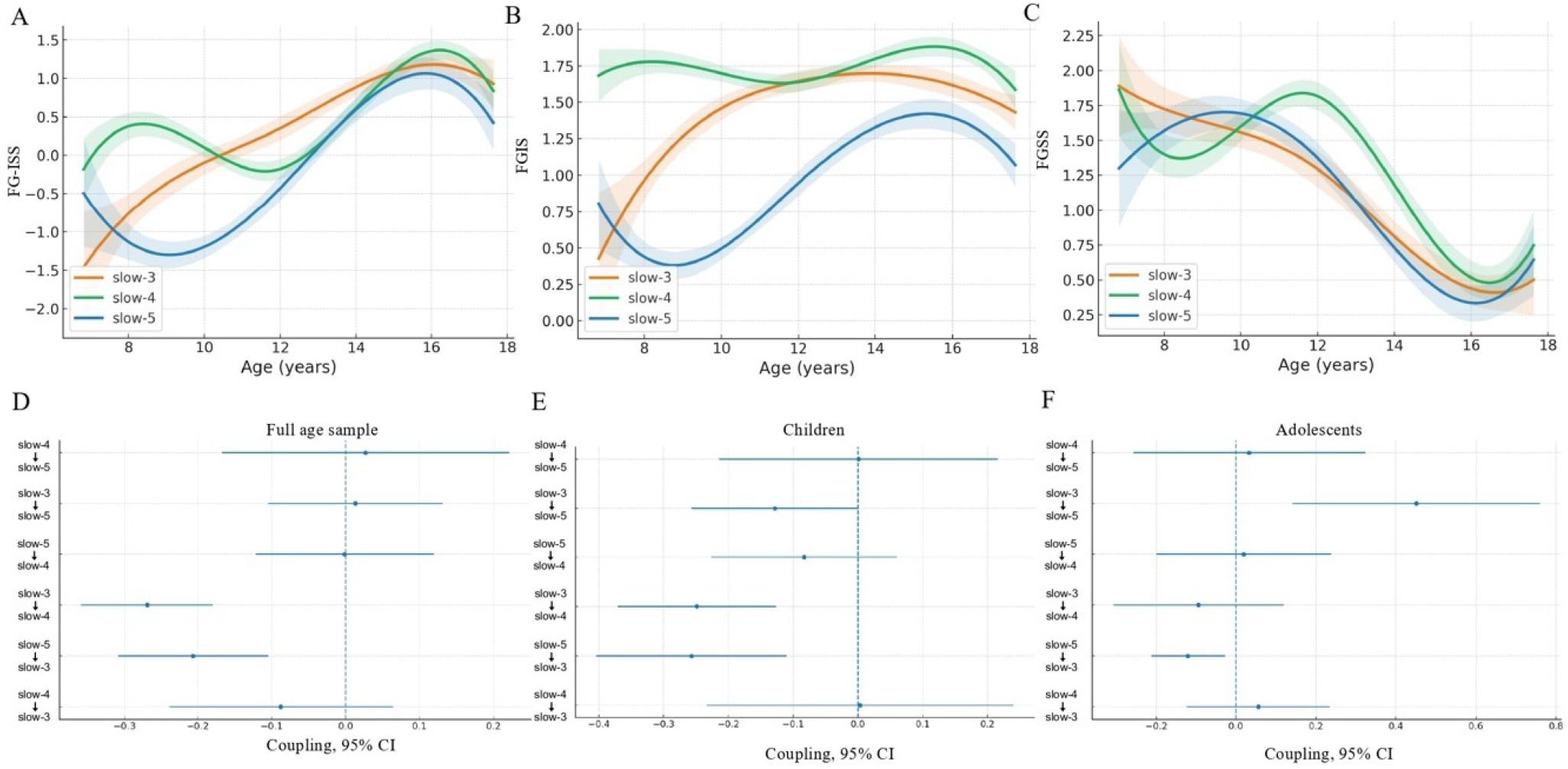
(A–C) Age-related growth curves for FG-ISS, FGIS, and FGSS across the slow-3, slow-4, and slow-5 bands. FG-ISS (the first-gradient integration-segregation score) indexes the balance of integration vs. segregation in the first gradient: positive values (>0) indicate an integration-dominant configuration (Stage III); values near 0 reflect a mixed/transitional configuration (Stage II); negative values (<0) indicate a segregation-dominant configuration (Stage I). FGIS (the first-gradient integration score) quantifies first-gradient integration (higher = greater integration). FGSS (the first-gradient segregation score) quantifies first-gradient sensorimotor segregation (higher = greater segregation). (D–F) Directed cross-band coupling of FG-ISS over development (latent change score models; see Methods). The y-axis lists band pairs; A→B denotes the effect of earlier FG-ISS in band A on the subsequent annualized change in band B. The x-axis shows the coupling coefficient (γ): negative values mean higher early A predicts slower growth in B; positive values mean higher early A predicts faster growth in B. (D) Full sample: two significant negative couplings (slow-3→slow-4 and slow-5→slow-3; FDR-corrected). (E) Children (≤12 years): the same two significant negative couplings (FDR-corrected). (F) Adolescents (≥13 years): one significant positive coupling (slow-3→slow-5) and one significant negative coupling (slow-5→slow-3; both FDR-corrected).

### Cross-band coupling of developmental changes

To probe how frequency bands influence one another over time, we applied a latent change score (LCS) model to test directional, cross-band coupling among FG-ISS trajectories. For each band pair, we asked whether earlier levels in one band predicted the subsequent annualized change in the other, controlling for age. In the full sample (Fig. 3D), two couplings survived FDR correction: slow-3 → slow-4 (γ = −0.2692, 95% CI [−0.3588, −0.1795], P<0.0001, q_FDR_<0.0001; R^2^ = 0.5177), such that higher early slow-3 FG-ISS predicted smaller subsequent growth in slow-4; and slow-5 → slow-3 (γ = −0.2062, 95% CI [−0.3081, −0.1043], P = 0.0001, q_FDR_ = 0.0002; R^2^ = 0.5961), indicating that higher early slow-5 FG-ISS predicted smaller subsequent growth in slow-3.

Age-stratified LCS models (Fig. 3E–F) localized these effects. In children (≤12 years), we replicated the two negative couplings: slow-3 → slow-4 (γ = −0.2488, 95% CI [−0.3711, −0.1265], P = 0.0001, q_FDR_ = 0.0004; R^2^ = 0.5871) and slow-5 → slow-3 (γ = −0.2571, 95% CI [−0.4042, −0.1100], P = 0.0006, q_FDR_ = 0.0018; R^2^ = 0.6115). In adolescents (>12 years), the coupling pattern shifted to a bidirectional interaction between slow-3 and slow-5: slow-3 → slow-5 was positive (γ = 0.4509, 95% CI [0.1422, 0.7595], P = 0.0042, q_FDR_ = 0.0252; R^2^ = 0.6429), while slow-5 → slow-3 remained negative but attenuated (γ = −0.1197, 95% CI [−0.2123, −0.0270], P = 0.0113, q_FDR_ = 0.0340; R^2^ = 0.7019). In sum, childhood cross-band coupling is dominated by negative influences (higher early organization in one band forecasts slower growth in another), whereas in adolescence the system reorganizes toward a reciprocal slow-3/slow-5 interplay in which more advanced slow-3 organization accelerates subsequent slow-5 change, and slow-5’s dampening effect on slow-3 weakens. This developmental shift dovetails with our stage-based results, highlighting evolving coordination among frequency-specific processes as the cortex transitions from segregation-dominant to integration-dominant organization.

### Accelerated maturation in children with elevated social sensitivity

To further examine how differences in maturation across bands relate to social and cognitive development, we next stratified the sample by individual differences in social sensitivity and in intellectual ability, allowing us to test whether multi-band developmental patterns, and their timing, diverge across groups defined by social or cognitive functioning. To index social functioning, we used the Social Anxiety Scale for Children (SASC) to capture each child’s sensitivity to social evaluation and stress (La Greca et al., 1988). Because this measure is validated up to age 17, only participants aged 6-17 contributed SASC scores. As shown in Fig. 4A, preadolescent children in our sample reported generally low social anxiety (group mean ≈ 4, SD ≈ 3). After the onset of puberty, scores increased markedly, especially from age 13 onward, such that group means exceeded 7 by mid-adolescence. Based on this distribution, we classified children with scores ≥7 within their age group as high in social sensitivity. Importantly, our cohort consists of typically developing children; thus, “high social sensitivity” here denotes relatively elevated sensitivity within the normative range, defined with respect to the age-specific mean, and its interpretation differs slightly between childhood and adolescence. Specifically, the ≥7 cutoff corresponds to roughly one standard deviation above the mean in preadolescence (i.e., children experiencing higher social anxiety than most same-age peers), whereas in adolescence it approximates an above-average level (i.e., the upper half of the distribution). We then examined connectivity-gradient development across the three frequency bands within this high-sensitivity subgroup.

**Fig. 4.**
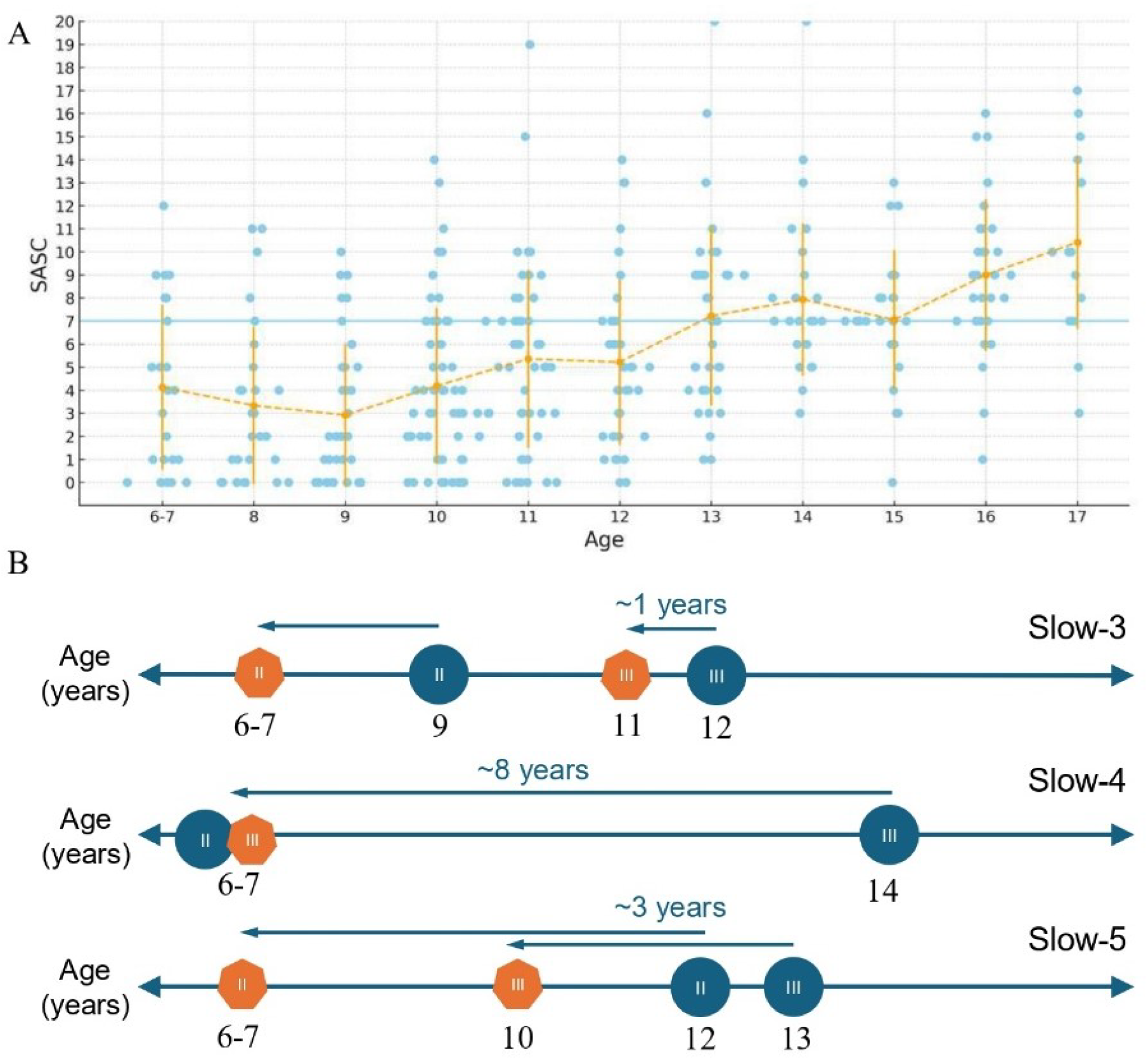
(A) Distribution of social anxiety scores across age groups. The x-axis indicates age groups, and the y-axis represents children’s social anxiety scores. A horizontal line marks the threshold at score = 7; within each age group, children scoring ≥7 were classified as the high social-sensitivity subgroup. (B) Earlier transition points into Stages II and III of Gradient 1 in the high social-anxiety group compared with the full sample, across all three frequency bands. Blue circles denote stage transition ages in the full sample; orange polygons indicate transition ages in the high social-sensitivity subgroup.

Relative to the full-cohort baseline, children with elevated social sensitivity attained adult-like gradient organization markedly earlier across all frequency bands (Fig. 4B, Fig. S4–S6). The canonical three-stage trajectory observed in the overall sample collapsed into a two-stage sequence in this subgroup, with the child-like Stage I essentially bypassed and an immediate entry into the transitional configuration. This leftward shift was frequency dependent.

By 6–7 years, the slow-4 band had already reached the stable Stage III pattern—an ∼8-year acceleration relative to the baseline onset at 14 years—whereas the slow-3 and slow-5 bands exhibited the mixed Stage II configuration at the same age. Stage III emerged at ∼11 years in slow-3 (≈1 years earlier than the cohort norm of 12 years) and at ∼10 years in slow-5 (≈3 years earlier than the 13-year norm). Thus, in every band the high-sensitivity group achieved mature gradient architecture before adolescence, with the largest advance in slow-4. Shifts of this magnitude, on the order of 1–8 years, indicate that heightened social sensitivity is accompanied by precocious large-scale cortical reorganization.

### Cognitive ability reflects frequency-specific maturation of functional gradients

To investigate the influence of general cognitive ability, we stratified participants by intellectual level using age-appropriate Wechsler IQ scores. As shown in Fig. 5A, IQ scores increased only gradually with age in our sample, especially compared to the steeper age-related rise observed in social anxiety scores. We focused on children ≤ 16 years (the upper age for WISC-IV norms) and first excluded any individuals with high social anxiety (scores > 1 SD above the age mean) to minimize confounding between social and cognitive factors. We then grouped the remaining children into three IQ brackets: a high-IQ group (Full Scale IQ ≥ 120), a mid-IQ group (100 < IQ < 120), and a low-IQ group (FSIQ ≤ 100). We examined the developmental trajectory of the first functional connectivity gradient across slow-3, slow-4, and slow-5 within each IQ group, comparing each to the full-sample baseline.

**Fig. 5.**
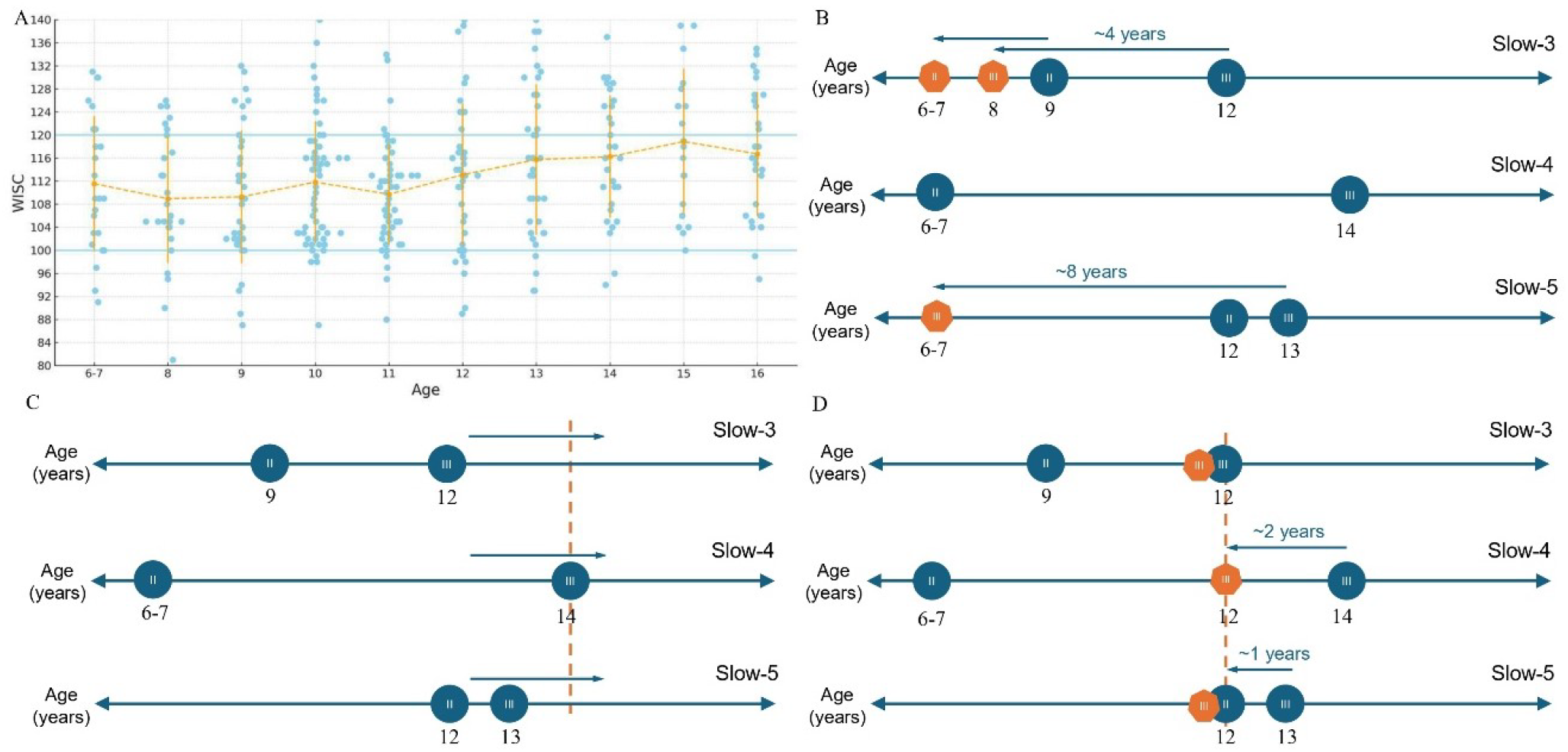
(A) Distribution of intelligence scores across age groups. The x-axis represents age groups, and the y-axis indicates full-scale IQ scores. Horizontal lines mark FSIQ = 100 and FSIQ = 120; within each age group, children were classified as high-IQ (FSIQ ≥ 120), mid-IQ (100 < FSIQ < 120), or low-IQ (FSIQ ≤ 100). (B–D) Developmental transition points into Stages II and III of the frist gradient across three frequency bands for children in the high-IQ (B), low-IQ (C), and mid-IQ (D) subgroups, compared with the full-sample baseline. Blue circles denote stage transition ages in the full sample; orange polygons represent transition ages in each IQ subgroup. In (B), high-IQ children show accelerated maturation in the slow-3 and slow-5 bands. In (C), low-IQ children show delayed development across all three bands, with none reaching Stage III by age 14. In (D), mid-IQ children exhibit synchronized Stage III onset at age 12 across all frequency bands.

Children in the high-IQ group showed earlier maturation of Gradient 1 in two of the three frequency bands (Fig. 5B, Fig. S7-S9). In slow-3, the high-IQ group reached Stage III by ∼8 years old (versus ∼12 years in the full sample). In the slow-5 band, Stage III was reached earliest, and by the widest margin, at roughly 6-7 years, an advance of about eight years relative to the ∼13-year baseline. By contrast, the slow-4 band showed no clear acceleration or delay in the high-IQ group’s trajectory relative to the norm, and its gradient pattern displayed no stable stage demarcation across the entire school-age period, possibly reflecting heterogeneous, network-specific developmental advantages within the modular circuits that dominate this frequency range. For the low-IQ group, sample size constraints required using broader age bins (6–8, 9–11, 12–14 years) to ensure stable gradient estimates, and children older than 14 were not analyzed in this group due to low sample number. Strikingly, across all three bands the low-IQ group did not reach Stage III in any age bin, indicating a generalized lag or under-development of the principal gradient compared to typical age expectations (Fig. 5C, Fig S10–S12). In the mid-IQ group, gradient maturation proceeded on an essentially normative timetable in all bands, with Stage III consistently attained by ∼12 years across slow-3, slow-4, and slow-5 (Fig. 5D, Fig. S13). This convergence around age 12 (in children with neither exceptional IQ nor high social sensitivity) suggests that ∼12 years may represent a normative milestone for multi-band cortical functional maturation when cognitive and social profiles are near the population average.

The band-specific patterns in the high-IQ group hint that superior cognitive ability is linked to the accelerated development of certain neural oscillation regimes. Notably, the two bands that showed early maturation in high-IQ children—slow-3 and slow-5—both correspond to integration-heavy functional levels in our framework. Slow-3 is associated with the perceptual integration component of the detection level, supporting the synthesis of sensory inputs and aspects of language processing. Slow-5 belongs to the modulation level, implicated in higher-order cognitive integration, self-generated thought, and multimodal association. Both bands thus facilitate integrative neural operations: one at the level of combining sensory information, the other at the level of unifying distributed association networks. Our findings suggest that earlier and more well-organized development of these integrative functional gradients may underlie superior cognitive performance. This is consistent with models positing that intelligence depends on efficient information integration across brain regions (Basten et al., 2015; Jung & Haier, 2007). Indeed, prior graph-theoretical studies have found greater global efficiency and more optimized network topology in youth with higher IQ (Suprano et al., 2019), and task-based fMRI studies link higher IQ with stronger connectivity in frontoparietal “executive” networks and the DMN (Sherman et al., 2014). Our results extend these findings by showing that high-IQ children not only have different connectivity strength at a given age but actually reach the mature stage of large-scale network integration earlier in development – at least for the frequency bands that support integrative functions.

In contrast, we saw no acceleration (and no clear advantage) in the slow-4 band for high-IQ children. Slow-4 primarily supports segregated, modular processing for core cognitive operations (the “computation” level). The non-stable acceleration in slow-4’s timeline may indicate that basic modular cognitive circuits mature on heterogeneous schedules and have long-lasting phases of developmental plasticity in gifted children, and that higher intelligence is more specifically related to integration across systems rather than the isolated development of any single cognitive module. Finally, the low-IQ group’s delayed trajectories across all bands suggest a pervasive lag in developing the brain’s functional hierarchy. Their principal gradient failed to attain an adult-like state even by mid-adolescence, which could reflect neurodevelopmental factors limiting the integration of information across both low- and high-frequency channels. When considered together, the subgroup analyses revealed a noteworthy convergence: in the absence of phenotypes that deviate from the mean (neither high nor low IQ, and no high social sensitivity), children tended to reach Stage III of gradient development around 12 years of age in all bands. This consistent ∼12-year transition in the “average” subgroup reinforces the idea that early adolescence is a typical inflection point for reorganization of cortical connectivity gradients. By contrast, both high social sensitivity and high or low IQ were associated with deviations from this normative timeline, either accelerating or delaying specific aspects of multi-band network maturation.

### Reproducibility of the three-stage developmental pattern

To assess the reproducibility of the multi-band functional connectivity gradient trajectory, we conducted a split-half analysis: the full cohort was divided into two subgroups, and the same age-binned, multi-band gradient pipeline was applied to each subgroup. Because behavioral factors can influence maturation rates, we used a nearest-neighbor pair-matching procedure to ensure that, within each age bin, the distributions of social anxiety and IQ were closely matched between the two subgroups. The resulting stage assignments are shown in Fig. S14. Overall, both groups reproduced the three-stage developmental pattern across frequency bands, and the onset of Stage III closely matched the full-sample estimates. In subgroup A, Stage II began ∼1 year earlier than in the full sample for slow-3 and slow-5; in subgroup B, the advance in these bands was somewhat larger. By contrast, slow-4 exhibited stable staging in both subgroups. Taken together, the split-half results confirm the reliability of the three-stage pattern and the stability of Stage III across bands and samples. The modest shifts in Stage I/II boundaries for slow-3 and slow-5 suggest greater sensitivity to inter-individual differences in social and cognitive measures during childhood, highlighting the need for larger samples or reduced heterogeneity in younger age bins to obtain maximally stable estimates.

## Discussion

Here we introduce a frequency-resolved view of developmental connectivity gradients that reveals features obscured by broadband fMRI. By decomposing BOLD activity into slow-3, slow-4, and slow-5 bands, we uncover a reproducible three-stage sequence of cortical gradient maturation whose timing is frequency specific. We further chart age trajectories capturing the balance between integration and segregation in the first gradient across bands and show that cross-band coupling reorganizes from predominantly negative influences in childhood to a bidirectional interplay between slow-3 and slow-5 in adolescence. Stratified analyses link these frequency-dependent timetables to meaningful individual differences: earlier, band-specific maturation in children with higher social sensitivity and higher IQ, and prolonged maturation in those with lower IQ, which underscores behavioral relevance. Together, these advances sharpen the mechanistic account of how distinct oscillatory timescales sculpt the maturing connectome and provide frequency-specific norms for future developmental and clinical biomarker studies.

### Multi-band analysis reveals frequency-dependent trajectories of cortical gradient maturation

Functional connectivity gradients capture the large-scale axes of cortical organization, with Gradient 1 typically spanning sensory–transmodal divisions and Gradient 2 distinguishing visual from somatomotor systems. These gradients provide a low-dimensional map of how functional networks are arranged, and their developmental reorganization reflects the consolidation of cortical hierarchy. Our findings show that the broad reorganization of functional brain architecture from childhood to adolescence is conserved across frequency bands, yet the developmental trajectories of connectivity gradients are strongly frequency-dependent. Consistent with earlier single-band work, we observe the expected adolescent “flip” of the first two gradients, wherein the spatial organization of Gradient 1 and Gradient 2 reverses as the cortical hierarchy consolidates (Dong et al., 2021; Xia et al., 2022). By fractionating the BOLD signal into narrower bands, however, we uncover a more nuanced three-stage maturation sequence and clearly delineate the intermediate transitional stage that bridges child-like and adult-like configurations, an effect only faintly apparent in broadband analyses (Dong et al., 2024). Notably, although multi-band studies in adults report largely consistent spatial patterns of the first two gradients across slow-3, slow-4, and slow-5 (Gong & Zuo, 2023), our developmental data reveal marked differences in pacing across bands, indicating that frequency-specific neural processes mature on distinct timelines.

A coherent ordering emerges across frequencies. Slow-3 matures earliest: both gradients reach adult-like organization before the lower-frequency bands, consistent with the idea that frequency ranges linked to more primary sensory-perceptual functions reach maturity sooner than those subserving higher-order association (Kawashima et al., 1995). In line with prior evidence implicating slow-3 in early sensory integration and semantic aspects of language (Frühholz et al., 2020; Gong & Zuo, 2024; Zhou et al., 2018), we observe that Gradient 1 undergoes the key transitional reconfiguration around ages ∼9–11, aligning with late childhood, a period marked by emerging integration of bottom-up sensory information and significant gains in linguistic skills such as dialogue and writing (Smith, 2005), and is adult-like by ∼12. Slow-4 shows the earliest onset but the most protracted course: Gradient 1 enters Stage II by ∼6– 7 years yet does not stabilize until ∼14, and Gradient 2 remains in the transitional state through much of school age, achieving adult-like organization only around ∼18–19. This extended timetable accords with the gradual elaboration of foundational cognitive and motor capacities across adolescence (Supekar et al., 2009). Slow-5, by contrast, exhibits a compressed peri-pubertal shift: both gradients reorient within a narrow window (∼11–13 years) and are adult-like by ∼13, consistent with this band’s centrality to DMN-related, internally oriented processes such as autobiographical memory and self-referential thought (Baria et al., 2013; Gohel & Biswal, 2015; Zhang et al., 2015). The timing aligns with the average onset of puberty and key milestones in identity formation (Waterman, 1982). Importantly, the slow-5 timetable closely matches the “flip” reported in conventional broadband (0.01–0.1 Hz) studies, suggesting that earlier single-band characterizations were dominated by slow-5 dynamics, whereas the slower-maturing slow-4 contributions were largely masked in the aggregate signal.

Taken together, fractionating BOLD into multiple bands demonstrates that each frequency range contributes uniquely to the temporal progression of cortical network maturation. At the group level, frequency-specific oscillations display distinct rates and reconfiguration patterns that map onto their putative functional roles within the cortical hierarchy, an organization that resonates with electrophysiological evidence that different oscillatory ranges support different facets of neurodevelopment (Bernardo et al., 2024; Khan et al., 2018; Marek et al., 2018; Schäfer et al., 2014; Wilkinson et al., 2024). These results argue against a single, broadband view of development and motivate a multi-band perspective for characterizing the mechanisms by which faster, more local oscillations and slower, integrative rhythms jointly sculpt cortical organization across childhood and adolescence.

### Frequency-specific developmental patterns and normative brain maturation

From a neurodevelopmental standpoint, the frequency-dependent trajectories observed in our study likely reflect hierarchical functional roles attributed to different frequency bands. The higher-frequency BOLD oscillations (slow-3) exhibited the earliest maturation, paralleling the early developmental timeline of primary sensory and perceptual functions and their corresponding cortical regions in childhood (A. C. Luo et al., 2024; Sydnor et al., 2023). In contrast, the slower oscillations (slow-5), associated with integrative and internally oriented cognitive processes, underwent rapid maturation around puberty, coinciding with the emergence of abstract thinking, self-identity, and enhanced self-referential capacities characteristic of early adolescence (Blakemore & Mills, 2014; Steinberg, 2005; Sweatman et al., 2025; van der Cruijsen et al., 2023; Waterman, 1982). Meanwhile, the intermediate slow-4 band spanned a more extended developmental window, suggesting that neural circuits supporting fundamental cognitive and motor functions undergo gradual refinement throughout the school-age years. This trajectory aligns with continuous developmental gains in attention regulation, academic proficiency, and motor coordination documented in this age range (Fosco et al., 2019; Mattison et al., 2023; Supekar et al., 2009; Volkmer et al., 2022; Zhang et al., 2025).

This frequency-specific partitioning aligns with the broader temporal hierarchy of brain maturation: foundational sensory systems develop first, providing a scaffold for higher-order cognitive, executive, and socio-emotional systems (Gogtay et al., 2004). Our multi-band results add nuance, showing that each hierarchical level may be preferentially supported by oscillations in distinct frequency ranges. Converging evidence across studies on different modalities shows that higher-frequency oscillations support lower-level, primary processing: they are more spatially local, reliably expressed in primary sensory and motor cortices, and strongly driven by bottom-up input (Frühholz et al., 2020; Gong & Zuo, 2024; Ray & Maunsell, 2011; Ryun et al., 2017). In contrast, lower-frequency oscillations are better suited for long-range coordination under conduction delays, enabling network-level integration and expressing robustly in attention, working memory, cognitive control, and canonical resting-state networks (Ma et al., 2021; Sadaghiani & Kleinschmidt, 2016; Vidaurre et al., 2018; von Stein & Sarnthein, 2000). Our study adds a developmental lens to this frequency-function hierarchy: bands associated with different functional hierarchical levels follow distinct maturational trajectories and show differential interactions with individual characteristics. Together, these findings argue that normative charts of brain development should adopt a multi-band framework to capture the full dynamics of hierarchical functional organization.

Identifying typical developmental trajectories and key timepoints is essential for understanding underlying neurodevelopmental mechanisms. There were notable differences related to social sensitivity and cognitive ability, highlighting the importance of variability for developmental benchmarks. Establishing normative developmental timelines also has clear clinical relevance (Hong et al., 2020). Deviations from the expected timing or sequence of gradient maturation, such as reduced adolescent changes or a prolonged Stage II, may signal impaired network integration. Evidence from autism spectrum disorder (ASD) provides an illustration: delayed and then aberrantly accelerated maturation of functional connectivity gradients is associated with persistent hierarchical imbalances in this condition (Lee et al., 2025). Multi-band analyses in clinical populations also reveal frequency-specific alterations in functional connectivity (Goodin et al., 2019; Makary et al., 2020; Martino et al., 2017; Russo et al., 2020; Su et al., 2021). Our findings provide a normative baseline of multi-band developmental trajectories against which clinical groups can be compared. By delineating frequency-specific developmental norms, this work offers a refined template for interpreting both individual variability and atypical patterns, highlighting how deviations from normative trajectories can inform understanding of neurodevelopmental health.

### Social sensitivity and precocious functional network maturation

Our analysis of the high social-sensitivity subgroup revealed markedly accelerated maturation of functional connectivity gradients across all frequency bands, with the greatest compression in slow-4. These children reached adult-like network organization years earlier than their peers, entering Stage III as early as 6–7 years and effectively compressing Stage II to the pre-school period. Heightened social sensitivity might accelerate the fine-tuning of social and emotional brain circuits, (Rudolph et al., 2021), aligning with the “stress-acceleration” framework: some children undergo faster development of emotion-related networks to meet social demands (Callaghan & Tottenham, 2016).

The intermediate Stage II in typical youth likely represents a window of heightened plasticity and network reorganization. High-sensitivity children bypass this stage, reaching adult-like patterns earlier, which may reduce flexibility but simultaneously enables early specialization for social monitoring. In particular, the slow-4 band, positioned between the high-frequency “detection level” and low-frequency “regulatory level”, serves as a critical window for integrating bottom-up and top-down processes, achieving maximal within-network integration (Gong & Zuo, 2025; Thompson & Fransson, 2015). Accelerated slow-4 maturation may enhance social processing, offering early adaptive advantages such as improved perspective-taking, but could limit broader experiential tuning (Miller et al., 2020; Tooley et al., 2021). By showing “adolescent-like” brain organization years before typical adolescence, these children illustrate that social brain development does not follow a single universal timetable, with temperament and environment shaping individual trajectories even within normative populations.

### Cognitive development and frequency-specific network integration

Children with higher IQs showed accelerated maturation of the principal connectivity gradient in the slow-3 and slow-5 bands, linked to local sensory integration and global high-order integration, respectively. The high-IQ subgroup reached Stage III earliest in slow-5, highlighting a link between superior cognitive ability and precocious development of distributed integrative networks. These findings support the Parieto-Frontal Integration Theory (P-FIT), which emphasizes efficient integration between distributed regions, particularly frontal and parietal cortices (Basten et al., 2015; Jung & Haier, 2007). Early maturation in slow-3 may facilitate rapid processing of external inputs, whereas early slow-5 integration supports abstract thinking, self-regulation, and memory (Aubry et al., 2021; Winner, 2000).

Interestingly, high-IQ children did not show accelerated maturation in the slow-4 band, where networks achieve maximal modular efficiency (Gohel & Biswal, 2015; Thompson & Fransson, 2015). This may indicate preserved plasticity in intermediate cognitive functions, contrasting with the high social-sensitivity group, whose precocity in slow-4 may come at the expense of flexibility. By contrast, lower-IQ children exhibited generalized delays across all bands, retaining child-like gradient patterns through mid-adolescence, suggesting delayed global functional maturation.

Overall, these results suggest that higher intelligence is linked to early integration across systems rather than accelerated maturation of mid-level circuits. This aligns with evidence that global efficiency and long-range connectivity, rather than modular efficiency, correlate with IQ (Basten et al., 2015; van den Heuvel et al., 2009; Langeslag et al., 2013). Our multi-band approach further clarifies the frequency context of these connections: fast oscillations (slow-3) support local sensorimotor integration, while slower oscillations (slow-5) underlie distributed association networks. Cognitive development thus reflects the coordinated integration of neural processes across multiple timescales, enabling the emergence of complex thought.

## Conclusion

We charted age-related reconfiguration of cortical connectivity gradients across multiple frequency bands, revealing previously unrecognized stages and individual differences in brain development. Typically developing children followed a reproducible three-stage trajectory, with each frequency band maturing according to its functional role. Individual differences in social sensitivity and intelligence demonstrate that brain development is not monolithic but modulated by both endogenous and exogenous factors. Our multi-band, multi-dimensional approach links oscillatory dynamics to cognitive and social outcomes, providing a framework to understand how intrinsic activity supports emerging mental abilities. Characterizing these frequency-specific developmental patterns may enable earlier detection of atypical maturation and guide interventions tailored to individual neurodevelopmental profiles.

## Methods

### Participants

We used neurodevelopmental data from school-age children collected in Chongqing, China, as part of the Chinese Color Nest Project (CCNP; Liu et al., 2021; Yang et al., 2017). Participants were recruited directly from schools. The CCNP adopted an accelerated longitudinal design, including twelve age groups ranging from 6 to 18 years at baseline. Each age group recruited eight boys and eight girls. Participants underwent one baseline assessment and two follow-up assessments, each separated by 15 months to control for seasonal effects. Initially, 192 children participated in the baseline assessment. After excluding subjects due to poor scan quality or dropout during follow-up, the project collected a total of 393 MRI datasets from 179 children. Participants were excluded from subsequent analyses if they had a history of neurological or psychiatric illness, a family history of psychiatric illness, organic brain pathology, a Child Behavior Checklist (CBCL) T-score higher than 70, or Wechsler Intelligence Scale for Children (WISC) standard scores below 80. Participants with excessive head motion (absolute framewise displacement > 0.5 mm) during resting-state scans were also excluded. Ultimately, 381 datasets were included in this study; age-group details and gender distributions are presented in Fig. 1A.

### Data Acquisition and Preprocessing

MRI data were collected at Southwest University using a Siemens Trio 3.0 Tesla scanner. Each session included two resting-state fMRI scans (each 7 min 45 s) separated by a structural T1-weighted scan (8 min 19 s). Only the resting-state scans were analyzed in this study. Resting-state scanning parameters were: repetition time (TR) = 2500 ms, echo time (TE) = 30 ms, flip angle = 80°, field of view (FOV) = 216 mm, matrix size = 72 × 72, 38 axial slices, slice thickness = 3 mm, inter-slice gap = 0.33 mm, in-plane resolution = 3 mm, and slice acquisition order = interleaved ascending (anterior-to-posterior phase encoding direction).

Resting-state data preprocessing followed the Connectome Computation System (CCS) pipeline (Xu et al., 2015). Steps included removal of the initial 10 s (4 TRs), head-motion correction, slice timing correction, de-spiking, estimation of head-motion parameters, registration of functional images to high-resolution T1 structural images via boundary-based registration, denoising using ICA-AROMA (Pruim et al., 2015), detrending, surface-based registration to cortical space (fsaverage5), and spatial smoothing with a 6 mm full-width at half-maximum (FWHM) Gaussian kernel.

### Multi-band Analysis

The preprocessed cortical surface-based time series data were further decomposed into multiple frequency bands using the DREAM toolkit (Gong et al., 2021). Based on BOLD signal sampling parameters (TR and scan duration), the DREAM toolkit utilizes the Natural Logarithmic Linear Law (N3L) frequency classification and the Nyquist sampling theorem to decompose the original BOLD fluctuations into distinct slow-wave frequency bands. Subsequent analyses were performed separately within each identified slow-wave band.

### Functional Connectivity Gradient Analysis

For each subject and frequency band, we first computed vertex-wise functional connectivity matrices (20,484 × 20,484) separately from the two resting-state scans in each session. These matrices were Fisher z-transformed and averaged within-subject, yielding one connectivity matrix per subject per frequency band. Next, group-level average connectivity matrices were computed within each age group, resulting in 12 age-group matrices per frequency band. Gradient analyses were conducted using the BrainSpace toolkit (Wael et al., 2020). Connectivity matrices were first sparsified, retaining only the top 10% of connectivity values per vertex, with remaining values set to zero. Cosine similarity was then calculated between each pair of vertices to generate a 20,484 × 20,484 affinity matrix. Dimensionality reduction of this affinity matrix was performed using diffusion map embedding, decomposing the data into principal eigenvectors representing the primary axes of variance. Each eigenvector defined a gradient across cortical regions.

### Behavioral measures

We assessed social functioning using the Social Anxiety Scale for Children (SASC) to index each child’s sensitivity to social evaluation and social anxiety (La Greca et al., 1988). The SASC comprises 10 items, yielding total scores from 0 to 20, with higher scores indicating greater social anxiety; its two-week test–retest reliability is 0.67. Children with SASC scores ≥ 7 within their age group were classified as high in social sensitivity. In childhood, this threshold approximates >1 SD above the age-group mean (i.e., relatively heightened social sensitivity among typically developing peers), whereas in adolescence it denotes scores above the age-group mean. We analyzed multi-band gradient maturation within this high-sensitivity subgroup to examine links between social sensitivity and developmental timing.

General cognitive ability was measured with the Wechsler Intelligence Scale for Children-Fourth Edition, Chinese version (WISC-IV), appropriate for ages 6–16. The WISC-IV (Chinese) shows strong psychometrics: split-half reliability >0.71 for subtests and 0.87–0.97 for composite indices; one-month test-retest reliability 0.71–0.86 for subtests and >0.80 for composites; and inter-rater reliability 0.96–0.99. Based on Full-Scale IQ (FSIQ), children in each age group were assigned to high-IQ (FSIQ ≥ 120), mid-IQ (100 < FSIQ < 120), or low-IQ (FSIQ ≤ 100) subgroups to evaluate how cognitive level relates to multi-band gradient maturation.

### Modeling age-related trajectories of multi-band first-gradient integration–segregation

During the school-age period, the first connectivity gradient (G1) typically progresses from a segregation-dominant organization toward an integration-dominant organization. To quantify this trajectory across frequency bands, we computed three metrics from each participant’s G1 map:

1. First-gradient integration score (FGIS). Let *gSMN*, *gDMN*, *gVN* denote the mean G1 coordinate of the somatomotor (SMN), default-mode (DMN), and visual (VN) networks, respectively. When G1 is dominated by the unimodal-to-transmodal integration axis (adult-like, Stage III), SMN and DMN lie at opposite ends of the axis and are far apart; when G1 is dominated by sensorimotor segregation (child-like, Stage I), DMN shifts toward the center and the SMN-DMN distance shrinks. We therefore define

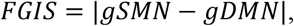

with larger values indexing stronger integration dominance.
2. First-gradient segregation score (FGSS). Under segregation dominance, SMN and VN anchor opposite ends of G1 and are far apart; under integration dominance they cluster toward the same end. We define

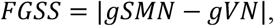

with larger values indexing stronger sensorimotor segregation.
3. First-gradient integration–segregation score (FG-ISS). A signed composite that summarizes the balance between integration and segregation:

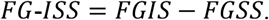

Positive values (FGIS > FGSS) indicate integration-dominant G1 (Stage III), values near zero indicate a mixed/transitional configuration (Stage II), and negative values indicate segregation-dominant G1 (Stage I).

Using 1–3 waves of longitudinal data per child, we modeled age-related trajectories of FG-ISS, FGIS, and FGSS in each band (slow-3, slow-4, slow-5) with generalized additive mixed models (GAMMs):

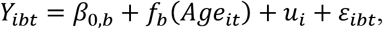

where *Y*_*ibt*_ ∈ {*FG* − *ISS, FGIS, FGSS*} for participant *i*, band *b* ∈ {*slow* − 3, *slow* − 4, *slow* − 5}, and time *t* ; *f*_*b*_ (*Age*_*it*_) is a band-specific smooth function of age (capturing nonlinearity); *u*_*i*_ is a subject random intercept; and *ε*_*ibt*_ is the residual.

### Cross-band developmental coupling

We next asked whether baseline organization in one frequency band predicts the subsequent annualized change in another band, after accounting for the target band’s own baseline and age. We used latent change score (LCS) model to estimate directional cross-band coupling among FG-ISS trajectories. For participant i, band b, and adjacent measurements t−1 → t, we defined the annualized change

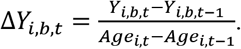

Coupling from band a to band b was tested via

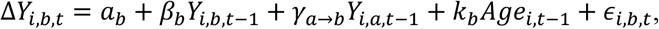

with the state update

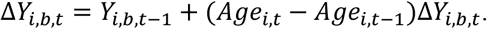

Here, *β*_*b*_ captures self-feedback (proportional change as a function of baseline in band *b*), *γ*_*a*→*b*_ is the directional cross-band coupling parameter of interest, and *k*_*b*_ adjusts for age at the earlier time point. We estimated all ordered band pairs among {*slow* − 3, *slow* − 4, *slow* − 5} and controlled the false discovery rate (FDR) across the set of coupling tests.

To localize the developmental window of any cross-band prediction, we repeated the LCS models without the age covariate, but stratified by baseline (transition start) age: Children (6 ≤ *Age*_*t*−1_ < 12 *years*) and adolescents (*Age*_*t*−1_ ≥ 13 *years*).

### Reproducibility validation

To assess the reproducibility of the three-stage developmental trajectory across frequency bands, we split the full cohort into two subgroups with matched distributions of social anxiety and intelligence scores. For each age group, we used a nearest-neighbor pair-matching procedure to ensure comparable score distributions between the two subgroups. Specifically, each participant’s social-anxiety and IQ scores were extracted and represented as a two-dimensional feature vector. Pairwise Euclidean distances were computed in this two-dimensional space, and the algorithm proceeded iteratively: starting from the first unassigned participant, the nearest unassigned neighbor (smallest Euclidean distance) was identified to form a matched pair. Within each pair, one participant was randomly assigned to Group A and the other to Group B, yielding balanced allocation while preserving the observed distributions of both measures within each age stratum. This procedure was repeated until all participants were assigned, resulting in two subgroups with closely matched baseline characteristics. After matching, we repeated the entire multi-band gradient pipeline separately in Groups A and B to evaluate the replicability of stage timing and cortical topography across independent, demographically comparable splits of the sample.

### Data Sharing and Open Science Practice

The CCNP datasets analyzed in this study are publicly available from Chinese Color Nest Data Community (https://ccndc.scidb.cn/en) at Science Data Bank.

## Notes

https://ccndc.scidb.cn/en

### Competing Interest Statement

The authors have declared no competing interest.

